# An open framework for biodiversity databases

**DOI:** 10.1101/010405

**Authors:** Miklós Bán, Sándor Bérces, Zsolt Végvári

## Abstract

OpenBioMaps is a recently developed, open and free web application with distributed database background, operated by several universities and national parks. This system provides free web map services and open database access by using standard OGC protocols. One of its’ main features is that users can create new and unique database projects. The databases involved are maintained by the data providers themselves. Standard tools supplied to the users include repeatable and referable data queries, exporting, evaluations and tracking of data changes. The system also provides a programmable data service for promoting data processing.

## 1. Introduction

Online accessible biodiversity-related databases, which are increasingly important for conservationists, are frequently used by the scientific community (Costello 2009). Although international agreements (10^th^ principle of the Rio Declaration 1992, Aarhus Convention 2001, Arzberger et al. 2004) considering public access to data and financial support for data sharing in conservation projects have already been drawn up, the number of freely accessible supraindividual biological databases is rather limited and the amount of publicly inaccessible biodiversity data is unexpectedly high (Reichmann et al. 2011). In contrast, open-access datasets can contribute substantially to the decrease of monitoring costs as an effectively cooperating network of citizen scientists collecting standardized biodiversity data can partly relieve the burden of surveying a number of taxa, many of which are often out of the scope of official or formal conservation goals (Hochachka et al. 2012). Governmental bodies, environmental companies, and green NGOs can benefit from freely available data using it in various GIS-based planning, for regional and rural development, and for linear infrastructure and industry, to provide sustainable development and management solutions especially when dealing with endangered or protected species.

The emergence of freely accessible datasets (e.g.: GBIF, Dryad, KNB, DataONE) opens the way for a number of unresolved issues in collection of biodiversity data. Firstly, open databases encourage users to start collecting long-term datasets, the inherent structure of which facilitates standardized data collection. Standardization of datasets can be a prerequisite for investigating a number of issues of key conservation importance: effects of global change on extinction probabilities, phenological shifts, behavioral adaptations, modification of community structure, ecological interactions, biological invasions, compilation of red lists, and the identification of ecosystem services which all might substantially influence conservation planning (Enke et al 2012). Secondly, the establishment of datasets with public access which consist of highly standardized data generates self-organized communities of citizen scientists, promoting community-level control of data collection methodology and facilitating data filtering to improve survey techniques and database quality (Bastow and Leonelli 2010, Magurran et al. 2010). Opponents of using open databases often argue (Enke et al. 2012) that (i) data collected by staff during work hours or using official technology belong to the particular institution and can be transferred to a third party only with a special permit, and (ii) data include spatially referenced records of species of key conservation value which can be easily accessed and, thus, potentially be harmful. However, note that (i) data acquired by using public money should be publicly accessible, and (ii) our dataset provides a unique possibility to develop multiple levels of data access rights allowing control of sensitive biodiversity data.

We present here a recently developed conceptualization and application (OpenBioMaps) of a free and open GIS-based database system specifically designed for scientific work and conservation management. This project was founded without government level control by universities and national parks, which is a basic difference compared to centralized systems such as GBIF (The Global Biodiversity Information Facility) or DataONE (Data Observation Network for Earth). This application provides a framework for structurally and functionally independent databases. The individually structured databases help to avoid overcomplicated structures created by generalization. The application serves as a long-term data repository and a common tool for data processing as well, considering the most common problems of small databases are long-term financial and technical support and maintenance (Bastow and Leonelli 2010). Furthermore it provides an individual level data storing tool (Whitlock et al. 2010) and a reliable sharing place (Savage and Vickers 2009) for individual projects. Our approach is to provide a reliable and simple tool which is applicable in everyday work for individual users, research teams and organizations alike. We facilitate data influx by simplifying the uploading process, therefore there is no centralized verification of data and no central restrictions while data transfer and also the database structures are as simple as possible. We also provide optional tools for different level data access, from the simple query to the data mining. This open approach promotes the usage of individual datasets of involved databases, however it requires more attention and unique solutions for processing data on higher lever.

## 2. System description

The OpenBioMaps system is a formal collaboration of institutes which was established to create a long-term operating, open database framework, which involves unique and independent biological datasets and provides free network services for data access and management.

The OpenBioMaps consortium was founded by University of Debrecen and Duna-Ipoly National Park Directorate and headed by a board with delegates from member institutes. This council maintains the system-level administrative tasks but does not govern the involved databases.

The system distinguishes between registered and unregistered users. Registered users (members) are entitled to access data in all functional databases and are able to invite new members without limitations and can create new, unique database projects. Special rights are provided for OpenBioMaps members founding a database, including the assignment of administrators, designing data field structure and the access rules.

Network services provided by the OpenBioMaps use OGC (Open Geospatial Consortium)-compatible standard protocols (WMS, WFS, Postgis) and non-standard protocols as well. These services are accessible through the *http://openbiomaps.org* portal by web browsers, desktop GIS applications (like QGIS) or private applications using advanced programmable interface. This latter interface uses a standard tool (JSON) for data query through a non-standard protocol called Project Data Service (PDS).

The databases are designed to store a broad variety of biological data with spatio-temporal and taxonomical attributes.

## 3. System tools

The web application offers several tools for users which facilitates querying, editing, importing, exporting and qualifying data, management of datasets and meta-data. In the web application, the database managers can set up unique textual query filter forms for their database which facilitate effective data searching for the users. For performing spatial queries, beyond the usual simple draw tools the users can upload geospatial vector data files which contains polygon definitions. The completed queries (including spatial queries) can be saved by registered users while saved spatial or textual queries are repeatable by any users using its references. Data contents of the databases are editable for users. Registered users can import data from various data sources and can create templates for regular imports (e.g. GPS data). Data query results are exportable to various file formats. Public reputation system provided by the OpenBioMaps is designed for data quality control by the community. Integrated tools (e.g. species name validation using the Catalogue of Life database) also helps to control data quality. All changes in the database are automatically saved and listed allowing searchable data history for users.

Although databases are not directly connected to each-other, some system level functions and structures are available which allows the application of higher level queries and constructing of meta-databases.

## 3. Databases and data access

The OpenBioMaps databases can be designed for various purposes (scientific, conservation management and data archiving). Currently the system comprises five databases with about 500,000 georeferenced records of approximately 5000 species from various types of investigations. The largest is the Duna-Ipoly National Park Directorate’s biotic database. Most of the data were collected by handheld GPS devices.

The databases contain georeferred data which is visualizable as (WMS) map layers in any OGC compatible GIS application (e.g. the web interface or the QGIS). The data attributes (through WFS or PostGIS) of the the map layers are also available which is suitable for filtering visualized data. In the simplest case, in the web application a user can perform spatial query by drawing a polygon, line or point on the map or create textual filters on species names or other attributes of a database. Results of the web queries can be downloaded as .csv or .gpx files and saved by registered users. The system automatically assigns unique identifiers for the saved queries. These identifiers are publicly available and simply usable in web browsers (e.g.: ‘http://danubedata.org/?LQ=48@6s0epbp0qa5ldlve’) or through the PDS service.

The PDS responses for the queries in easily processable JSON (http://www.json.org/) or xml formats and this service is available for several type of queries, for example retrieve species list of a database (http://openbiomaps.org/pds/?service=PFS&table=danubefish&specieslist) or retrieve data from a previously performed and saved query using its reference identifier (http://openbiomaps.org/pds/?service=PFS&LQ=48@6s0epbp0qa5ldlve) or query the all occurrences of a species in a project (http://openbiomaps.org/pds/?service=PRS&table=danubefish&species=Abramis+brama&type=xml).

## 4. Discussion

OpenBioMaps is a **unique and free** framework for biological databases which offers a wide range of functionalities, including integrability. The involved projects have unique data structure, however the system level data integrity is implemented by general functions and meta-data structures. The data can even be accessed from within scientific tools such as the R statistical programming environment that greatly extends the range of applicability of the system. Further, data provided by OpenBioMaps can be viewed and edited using a wide range of spatial analyst applications such as QGIS that allows fast and easy management of spatial data (Urbano and Cagnacci 2014) leading to accelerated publication processes.

To implement a long-term sustainable project we designed OpenBioMaps by using only widely supported free softwares, and at the same time by ensuring low operation costs as maintenance roles were transferred to registered users. OpenBioMaps is a decentralized and distributed system operated by several universities and national parks. Therefore, our system offers a means of improved **cooperation** between conservation management and science, **citizen scientists** and conservation authorities. This might have multiple benefits for maintenance and research of biodiversity. Firstly, cooperation and exchange between science and conservation can contribute substantially to reach our shared objective of halting biodiversity loss within a decade. Secondly, effective cooperation can induce positive feedbacks in usage of public databases: the involvement of public knowledge on biodiversity promotes the increase the number of registered users of OpenBioMaps. Thirdly, growing knowledge in specific fields might intensify the focus on less investigated subjects, which facilitates the exploration of data-deficient taxa and habitats.

As all databases contain at least taxonomic, temporal and spatial information, **meta-analyses** of various datasets open the way towards a wide range of biodiversity analyses, similarly to museum and herbarium collections. For example, such meta-analyses allows the assessment of parameters of population dynamics on larger temporal and spatial scales.

The database framework presented here should contribute critically and significantly to the integration of individual databases into publicly funded research on a global scale. The new system advocated here also expands the horizon for environmental education, e.g. by involving students in wildlife data collection, databases usage and analyses with real data sets.

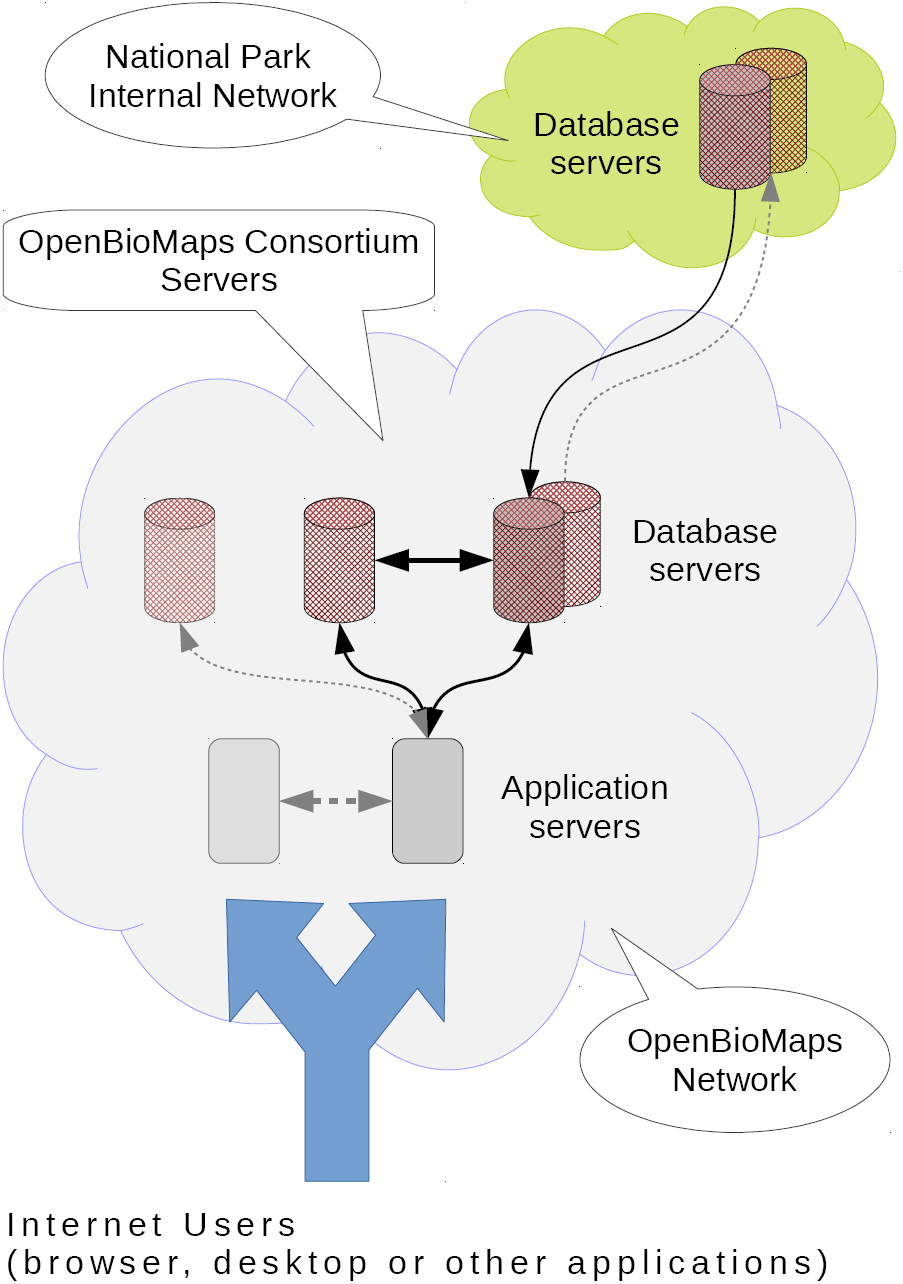
Internet Users (browser, desktop or other applications)

## Acknowledgments

This research was supported by the European Union and the State of Hungary, co-financed by the European Social Fund in the framework of TÁMOP-4.2.4.A/ 2-11/1-2012-0001 ‘ National Excellence Program’. We are grateful for Prof. Mark Hauber, Dr. Gábor Sramkó, Dr. Mihály Földvári and Dr. Jácint Tökölyi for improving the manuscript.

